# Computational mapping of the differentially expressed gene-lncRNA pairs present at the root nodule developmental stages of *Arachis hypogaea*

**DOI:** 10.1101/724674

**Authors:** Ahsan Z. Rizvi, Kalyani Dhusia

## Abstract

RNA-sequencing (RNA-seq) data analysis of the different stages of root nodules formation in peanut *Arachis hypogaea* investigate the genetic features. Genes related to the root nodules formations in this plant are extensively studied [1] [2] [3] [4] [5], but less information is present for their relations with long noncoding RNAs (lncRNAs). Bioinformatics techniques are utilised here to identify the novel lncRNAs present in the publically available RNA-seq data reported [6] for the different stages of root nodules formation in this plant. Highly correlated, significant, and Differentially Expressed (DE) gene-lncRNA pairs are also detected to understand the epigenetic control of lncRNA. These pairs are further differentiated between *cis* and *trans* antisense lncRNAs and lincRNAs based on their functions and positions from the genes. Obtained results are the catalogue for the highly correlated and significant DE gene-lncRNA pairs related to root nodules formation in *A. hypogaea*.

## 1 Introduction

Development of computational techniques of RNA-seq data analysis has increased our understanding for non-coding RNAs (ncRNAs). LncRNAs are the diverse class of ncRNAs with the length more than 200 nucleotides. These transcripts lack protein-coding potentials or have partial coding possibilities [7] [8]. LncRNAs are transcribed in plants by polymerase II, III, IV, V [9] and processed by splicing, non-splicing, and polyadenylation [10]. Classifications of lncRNAs are based on their locations related to genes, targeting mechanism, and functional mechanism. A subgroup of lncRNAs is long intergenic non-coding RNAs (lincRNAs) which are transcribed from sense strand and intergenic regions [11]. These are involve in stress response, signal transductions, and developmental process of the plants [12]. Few lncRNAs are transcribed from the antisense strand of the transcriptional units. These antisense lncRNA are involved in several cellular activities like transcription [13], translation [14], and *cis-trans* regulation [15] of genes. *Cis*-antisense lncRNAs act in nearby genes while *trans*-antisense lncRNAs act in any distant gene. Therefore, a correlation test [16] is required between the expressions patterns of genes and antisense lncRNAs to check the mutual influence.

LncRNAs show the response to external and internal stimuli [17] and have the role in the tissue development in the plants [18]. Genome-wide computational techniques are used for identification and functional analysis of lncRNA in the several plant species [3] [7] [19]. LncRNA plays a vital role in the stimuli responses in few plants like *Arabidopsis thaliana* [20]. Poly(A)+ and poly(A)- lncRNAs of this plant show active role in Pi-starvation [21]. Shorter lncRNA poly(A)- is more specific to stimuli and involved in the pathway related to Pi-starvation. Flowering time in *A. thaliana* is regulated by a floral repressor *FLOWERING LOCUS C (FLC)*, antisense lncRNA *COOLAIR*, and sense lncRNA *COLDAIR* [22]. These two types of lncRNAs play an important role in the epigenetic control of *FLC* via histone modifications. Photomorphogenesis in the same plant is also regulated by lncRNA *HIDDEN TREASURE 1 (HID1)* [10]. This lncRNA negatively governs the PIF3 gene by binding the promoter region.

Computational techniques are reported to the model legume species *Medicago truncatula* for the identification and classification of lncRNAs [23]. Cortical cells of the nodulated roots of *M. truncatula* show the expression of lncRNAs *MtENOD40* [24] [25] which codes two short polypeptides but lack long Open Reading Frames (ORFs). This lncRNA interacts with RNA-binding proteins *MtRBP1* and moves from the nuclear speckles to the cytoplasm during root nodulation process. LncRNA *GmENOD40* [26] of *Glycine max* has the similar functions for the root nodule formation. This lncRNA is expressed in the root nodule vascular bundle of the matured *G. max*. Induction of *GmENOD40* in the vascular bundle of the root nodules is required the presence of intracellular bacteria or infections. These studies demonstrate the importance of lncRNA for the root nodule formations in the leguminous plants.

Leguminous plants have symbiotic relationships with *Rhizobium* bacteria in their roots. These relationships play a significant role in Nitrogen fixation of the soil. Rhizobial invasion and root nodule organogenesis is the host controlled in the majority of the leguminous plans [25]. Rhizobia enter through infection thread and root nodule primordium in these plants. As an exception, rhizobia directly invade root cortical cell via epidermal cracks in dalbergoid legumes like *Arachis hypogaea* [6]. Wild diploids parents of this plant are *Arachis duranensis* and *Arachis ipaensis* and their genomes are used for mapping purpose. Transcriptome data are downloaded for the different stages of symbiosis in *A. hypogaea* like root invasion stage (1DPI), root nodule primordia formation (4DPI), the spread of infections in the root nodule (8DPI), immature root nodules with rod-shaped rhizobia (12DPI), and mature root nodules with spherical symbiosomes (NOD). Differential gene expression for the progress of symbiosis in *A. hypogaea* are reported [2] [3] [4] [5] [6] but their epigenetic connections with lncRNAs are explored in this study.

Patterns of the DE lncRNAs are correlated with the patterns of DE genes to understand the epigenetic role of lncRNAs. The biological significance of the highly correlated and significant DE gene-lncRNA pairs are explained in this studies. These pairs are further differentiated into gene-lincRNAs pairs, *cis*-antisense gene-lncRNA pairs, and *trans*-antisense gene-lncRNA pairs. Functional analysis of these pairs is also performed to understand the biological processes that could be influenced by lncRNAs.

## 2 Method

### 2.1 LncRNAs identification and validation

RNA-seq data are downloaded from https://www.ncbi.nlm.nih.gov/sra?term=SRP107173 [6]. These data contain transcriptomic expressions of six root nodule developmental stages after the infection with *Bradyrhizobium* Semia 6144. Three replicates per stage are converted into single ended “*.fastq” files through “fastqdump”. These files are passed through Trimmomatic *v*0.36 [27] for the removal of adapters and low-quality nucleotide bases. The trimmed reads are mapped with genomes of two diploid parents of *A. hypogaea* which are *A. duranensis* and *A. ipaensis* by HISAT2 [28]. Resulted BAM files are assembled by StringTie *v*1.3.2 [29]. Assembled gene transfer format (GTF) files are merged with “stringtie-merge” for each genome. Mapped transcripts are extracted by “gffread-w” with the inputs of respective merged GTF files and genome files. CD-HIT-EST [30] removes the redundancies of extracted transcripts, and the non-redundant transcripts are used as input for the possible lncRNAs identifications.

A “Python” [31] based pipeline is developed to filter the expected lncRNAs from the total mapped transcripts. First, it removed transcripts less than 200*bp* long and stored in a FASTA file. The second step is the removal of transcripts with ORFs more than 120 amino acids in length. In the third step, filtered transcripts are checked for homology with the known protein. Internal BLASTX [32] is used to align transcripts with universal protein (UniProt) consortium [33] database which is downloaded from (http://www.uniprot.org/downloads), dated 7^*th*^ March 2018. Transcripts align with proteins with e-value less than 0.001 are expelled from the transcripts database. This alignment check reduced the false positive scoring cases formed due to the translation of pseudo-genes. Filtered transcripts are uploaded on the online version of Coding Potential Calculator 2 (CPC2) [34] for the validation. This calculator uses Support Vector Machine (SVM) [35] over the four intrinsic parameters such as Fickett TESTCODE score [36], ORF length, ORF integrity, and Isoelectric point (pI). Another “Python” script is developed which takes the input of the results of CPC2 and removes “coding” and “partial coding” transcripts from the long noncoding transcripts database. Presences of transfer RNAs (tRNA) in the long noncoding transcripts database are checked by ARAGORN (http://mbio-serv2.mbioekol.lu.se/ARAGORN/index.html) [37]. Fast and sensitive SortMeRNA [38] is used to remove ribosomal RNA (rRNA) from the long noncoding transcripts database. These filtered long noncoding transcripts database are consider as potential lncRNAs.

### 2.2 Differential expression analysis of lncRNAs

Merged GTF files which are initially used for potential lncRNA identification is further utilised for reassembling of BAM files with “stringtie -eB -c 3”. Resulted files have the details of mapped transcripts with chromosomes numbers, start, end nucleotide position and their respective attributes. Attributes contain semicolon separated values of fragments per kilo-base of transcripts per million read pairs (FPKM) and transcripts per million (TPM) values for each mapped transcripts. Read counts for the mapped transcripts are generated by “prepDE.py” downloaded from (https://ccb.jhu.edu/software/stringtie/dl/prepDE.py). Chromosome number, nucleotide locations and FPKM values are identified for the potential lncRNAs. Significant FPKM values of the potential lncRNAs are further utilised for Log2 Fold Change (Log2FC) analysis. FPKM values at UI stage are considered as a control (C). While 1DPI, 4DPI, 8DPI, 12DPI, and a NOD are used as treatment (T) for the Log2FC calculation. Transcripts Log2FC ≥ 1.0 with P-value < 0.05 are consider as up-regulated. Similarly, Log2FC ≤ −0.05 and P-value value < 0.05 are consider as down-regulated. Similarly, a Log2FC data are generated for genes of the both genomes. DE lncRNAs are searched within the range of 5*kb* nucleotide position [39] [40] from the genes. If any significant lncRNA is observed within this range, then consider as gene-lncRNA pairs. Gene Ontology identity number (GOids), and KEGG Orthology identifier (KOids) of the paired genes are extracted from their respective Generic Feature Format (GFF) file. GO enrichment analysis [41] [43] is performed to check the biological activity of gene-lncRNA pairs. Log2FC values at the different stages of root nodulations of the paired genes and paired lncRNA are mutually correlated. This correlation is performed through Pearson correlation test [16], and significant and highly correlated gene-lncRNA pairs are identified. These pairs are classified into DE gene-lincRNAs pairs, *cis*-antisense gene-lncRNA pairs, and *trans*-antisense gene-lncRNA pairs based on lncRNAs locations from the gene [12] [42]. KOids related to these pairs are identified and utilised for the biological pathway analysis.

## 3 Results

Downloaded RNA-seq data contain 1494542967 reads with an average of 83030164 reads per samples. These RNA-seq sample files are stored in the compressed *.*fastq.tar.bz* format and passed through the quality check and adapter removal. Average 8.66% per samples are removed as adapters or low-quality nucleotide bases. Trimmed and compressed RNA-seq samples files are used for mapping along with *A. duranensis* and *A. ipaensis* genomes. RNA-seq samples files mapping with *A. duranensis* genome shows average 86.42% alignment rate while mapping with *A. ipaensis* genome shows average 86.65% alignment rate. Thirty-six *.*sam* files are resulted from the mapping process which are further converted into sorted, and indexed *.*sorted.bam* files. These files are used as inputs for assembling process by supplying the genomes annotations. Resulted *.*gtf* files are merged for each genome. Assembled transcripts are extracted from with their respective *.*merged.gtf* and genomes files. Non-redundant transcripts are further utilized for the identification of lncRNAs.

Total 33557 transcripts are filtered for *A. duranensis* while 37664 transcripts for *A. ipaensis* by the developed pipeline. Filtered long noncoding transcripts are uploaded on CPC2 (http://cpc2.cbi.pku.edu.cn) for the validation. The “coding” and “partial-coding” transcripts are removed from the long noncoding transcripts database by comparing with CPC2 results. Total 29522 transcripts are labelled as noncoding for *A. duranensis* genome while 33191 transcripts are labelled as noncoding for *A. ipaensis* genome by this calculator. Presence of rRNA and tRNA are checked in the long noncoding transcripts database, and total 32 tRNAs and 957 rRNAs are detected for *A. duranensis*. Similarly, 23 tRNAs and 995 rRNAs are detected for *A. ipaensis*. These tRNAs and rRNAs are removed from the long noncoding transcripts database, and remaining transcripts are considering as the potential lncRNAs. Chromosome number and nucleotide positions of the identified lncRNAs are obtained from their respective *.*merge.gtf* files.

It is observed that *A. duranensis* genome has 48.62% lncRNA found on “.” undefined strand, 25.69% on “-” antisense strand, and 25.68% on “+” sense strand while *A. ipaensis* genome contains 46.07% on “.” undefined strand, 27.31% on “-” antisense strand, and 26.61% on “+” sense strand. Fig. 1 and Fig. 2 are showing the distributions of lncRNAs over the chromosomes for both the genomes. Chromosome “A03” of *A. duranensis* shows highest counts for lncRNAs while “B09” of *A. ipaensis* genome has highest lncRNA counts. Distributions of lncRNA counts at the different length thresholds are calculated, and the results are given in Table 1. Average length 619*bp* is obtained for lncRNAs of *A. duranensis* while *A. ipaensis* has the average length 1357*bp* for lncRNAs. Similarly, the average length of mRNA is 1420*bp* for *A. duranensis* and 1357*bp* for *A. ipaensis*. These results show that the average length of lncRNAs is shorter than mRNAs for both the genomes. The average length of the ORFs are 14*aa* for lncRNAs and 27*aa* for mRNAs in both the genomes. These observations show that the average lengths of ORFs in mRNAs are higher than the average length of lncRNAs for both the genomes. Log2FC values at the different stages of root nodule formations are generated for genes and lncRNAs for both the genomes. Fig. 3 and Fig. 4 are showing the bar-plots for the up and down regulated lncRNAs obtained at the different stages of root nodule formations for both the genomes. Numbers of up-regulated lncRNAs are increasing from 1DPI to 8DPI and then start decreasing for both the genomes.

**Fig. 1.**
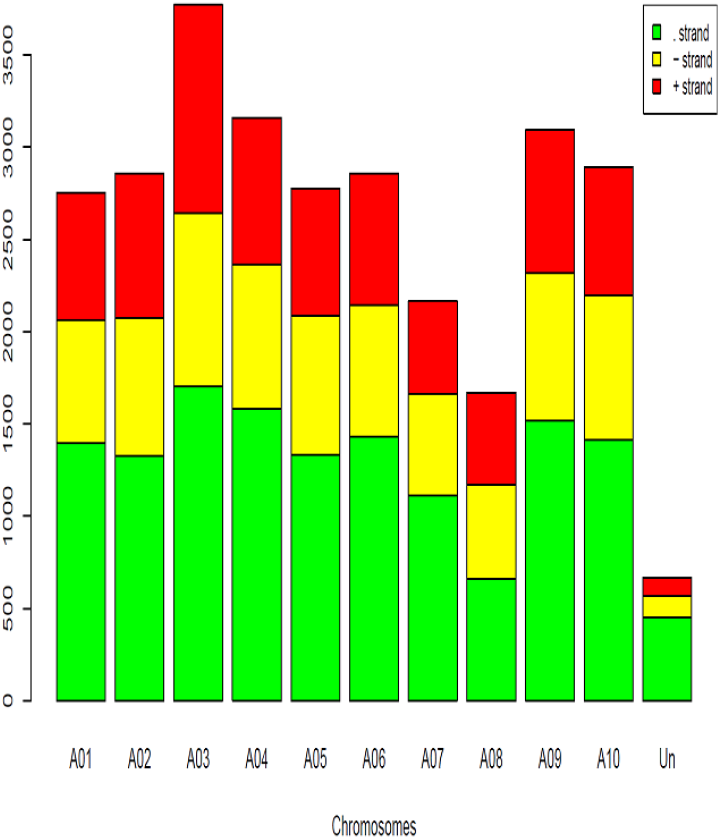
The distribution of IncRNAs over the chromosomes of *A. duranensis*.

**Fig. 2.**
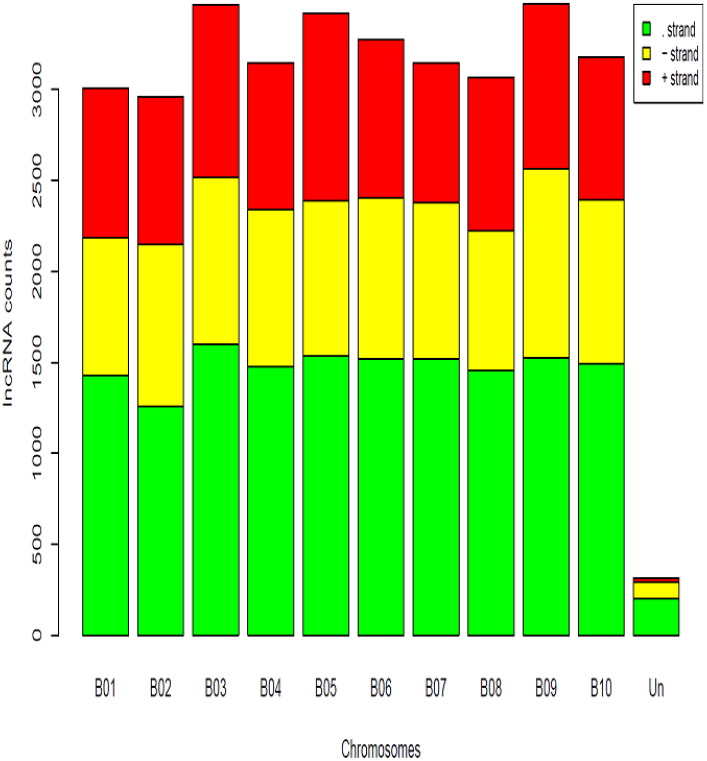
The distribution of lncRNAs over the chromosomes of *A. ipaensis*.

**TABLE 1.** LncRNAs and mRNAs counts at the different lengths ranges for *A. duranensis* and *A. ipaensis*.

| Length range | *A. duranensis* |  | *A. ipaensis* |  |
| --- | --- | --- | --- | --- |
|  | lncRNA count | mRNA count | lncRNA count | mRNA count |
| 0bp to 2000bp | 27839 | 28724 | 30989 | 33414 |
| 2001bp to 4000bp | 741 | 6985 | 1242 | 7342 |
| > 4000bp | 90 | 1025 | 120 | 1084 |

**TABLE 2.** Counts and classifications of GO terms of highly correlated and significant DE gene-lncRNA pairs of *A. duranensis* and *A. ipaensis*.

|  |  | LncRNAs of *A. duranensis* |  |  | LncRNAs of *A. ipaensis* |  |  |
| --- | --- | --- | --- | --- | --- | --- | --- |
|  |  | *cis*-antisense | *trans*-antisense | lincRNA | *cis*-antisense | *trans*-antisense | lincRNA |
| Total no. of genes |  | 18 | 47 | 203 | 29 | 67 | 194 |
| Annotated genes |  | 9 | 26 | 48 | 20 | 42 | 33 |
| GO terms | Biological process | 6 | 15 | 22 | 12 | 31 | 14 |
|  | Cellular component | 1 | 4 | 7 | 3 | 12 | 3 |
|  | Molecular function | 9 | 25 | 30 | 13 | 37 | 28 |
| Counts for $CC \geq 0.85$ | | 13 | 32 | 180 | 23 | 49 | 117 |
| Counts for $CC \leq -0.85$ | | 5 | 15 | 23 | 6 | 18 | 17 |

**Fig. 3.**
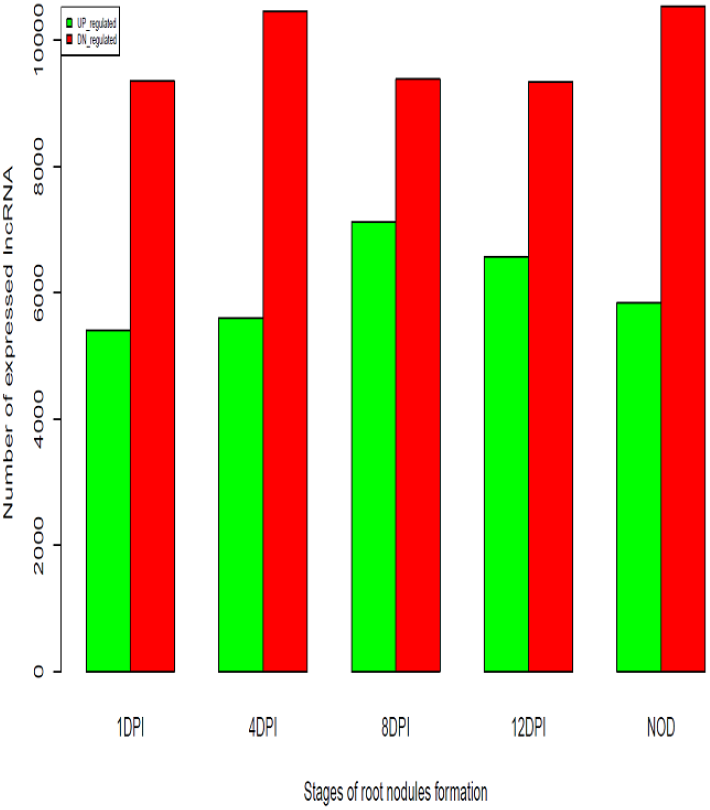
Total number of up and down regulated lncRNAs at the different stages of root nodule formation in *A. duranensis*.

**Fig. 4.**
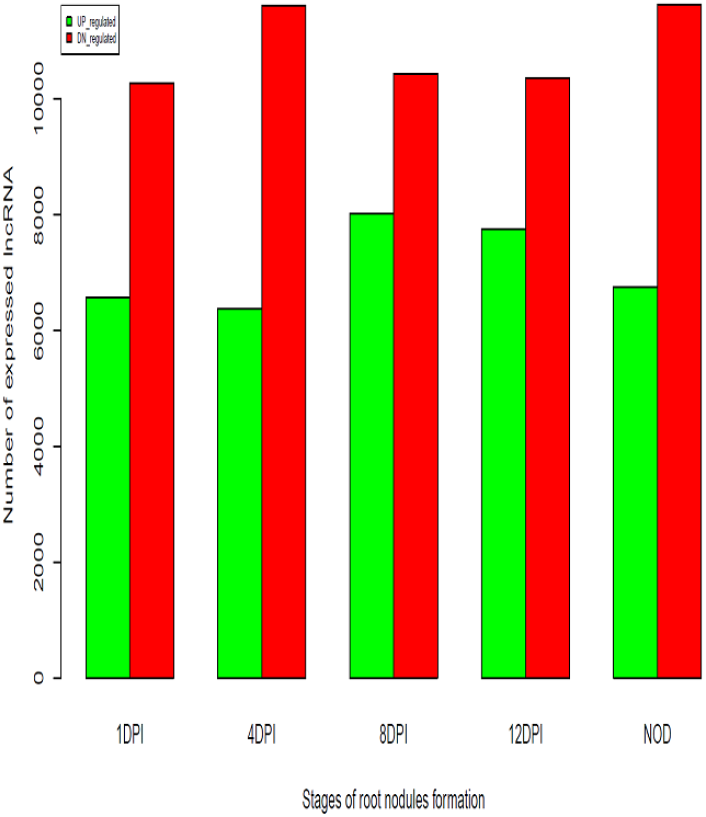
Total number of up and down regulated lncRNAs at the different stages of root nodule formation in *A. ipaensis*.

Nearest lncRNAs are searched within the range of 5*kb* nucleotide positions [39] [40] for each gene. Total 28670 gene-lncRNA pairs are identified for *A. duranensis*, and total 32351 gene-lncRNA pairs are identified for *A. ipaensis*. Fig. 5 is showing counts for the DE gene-lncRNA pairs at 1DPI, 4DPI, 8DPI, 12DPI, and NOD stages. Highest numbers of DE gene-lncRNA pairs are obtained at NOD stage for both the genomes. At this stage, total 3088 DE gene-lncRNA pairs are obtained for *A. duranensis* and 3470 for *A. ipaensis*. It is observed that 1653 pairs are present in the same (++ or - -) strands or sense strand of the genome while 1435 pairs are in the opposite (+ - or - +) strands or antisense strand for *A. duranensis*. Similarly, 1827 pairs are present in the sense strand while 1643 pairs are directed at the antisense strands for *A. ipaensis*. GOids are extracted from GFF annotation files for the genes present in the gene-lncRNA pairs for both the genomes. GO enrichment analyses are performed with the P-value < 0.05. This analysis gives enriched GOids for *A. duranensis*, which are listed in Table 3. Similarly, enriched GOids are obtained for *A. ipaensis*, which are listed in Table 4. This analysis gives 27 enriched GOids for *A. duranensis* and 32 for *A. ipaensis*. Fig. 6 is showing a bar-plot for the total number of up-regulated genes counts with GO definitions common for both the genomes. Gene ontology consortium http://www.geneontology.org/ explain the biological importance of these enriched GO ids in term of biological process, molecular functions, and cellular components.

**Fig. 5.**
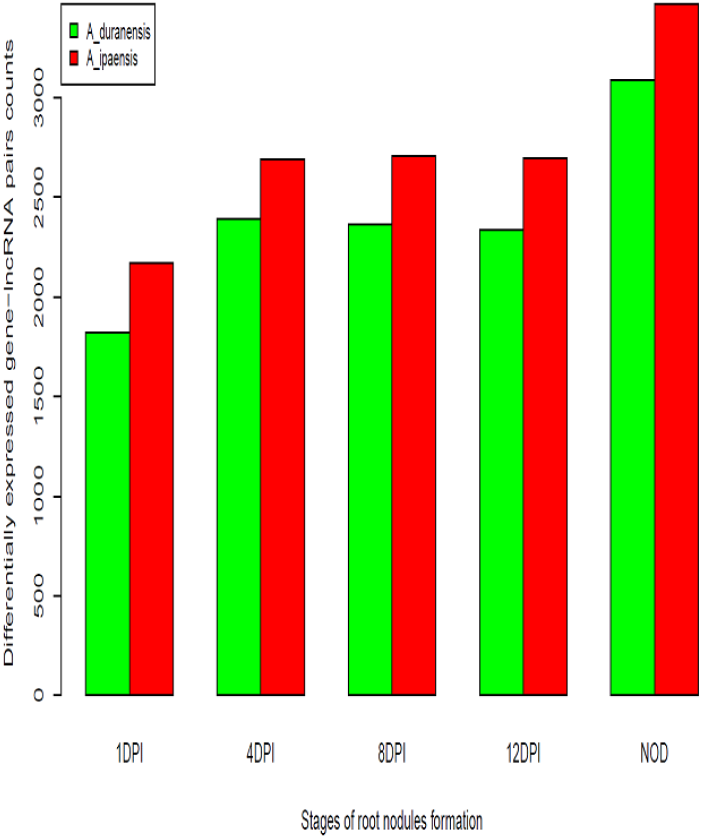
The total number of DE gene-lncRNAs pairs at the different stages of root nodule formation in *A. duranensis* and *A. ipaensis*.

**TABLE 3.**
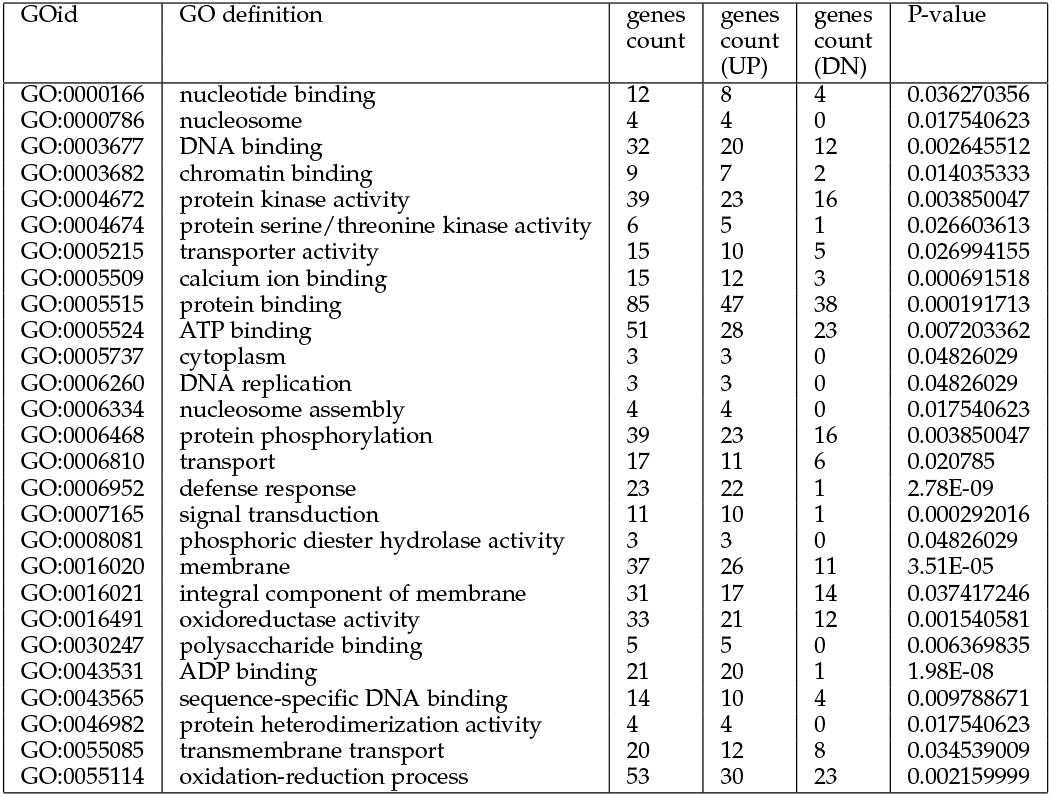
Enriched GOids of DE gene-lncRNAs pairs for *A. duranensis*.

**TABLE 4.**
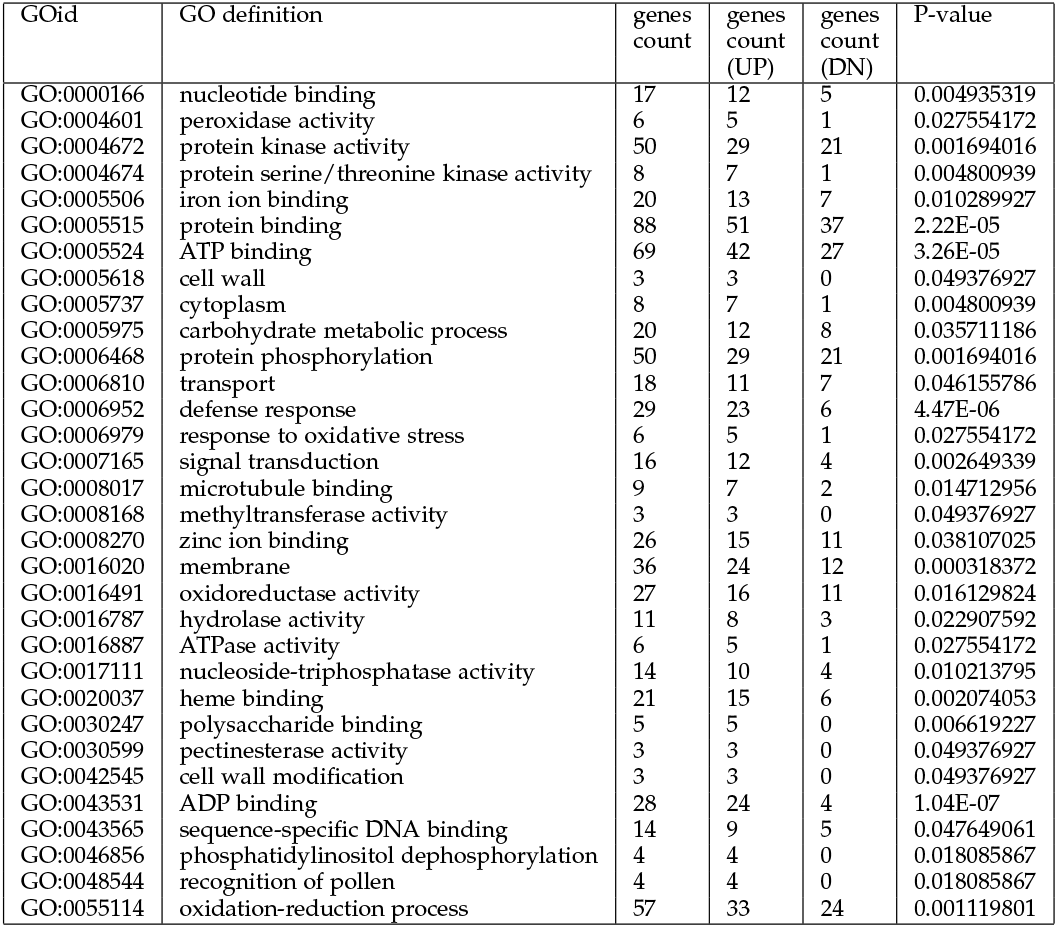
Enriched GOids of DE gene-lncRNAs pairs for *A. ipaensis*.

**Fig. 6.**
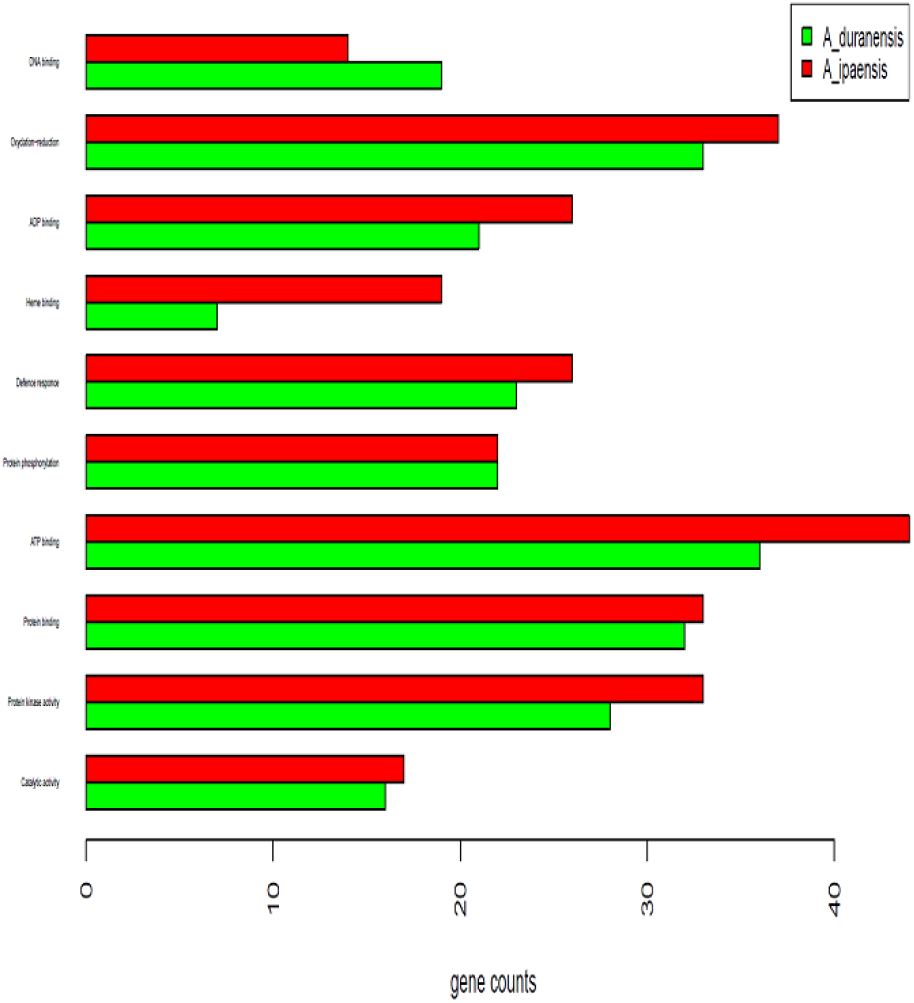
Up-regulated gene counts for the significantly expressed GO definitions common for *A. duranensis* and *A. ipaensis*.

Correlation tests are performed between the expression patterns of genes and lncRNA at the different stages of root nodule formations of *A. hypogaea*. Significant and highly correlated gene-lncRNA pairs are identified to understand epigentic control of lncRNAs. Table 5 is showing *cis*-antisense gene-lncRNA pairs for *A. duranensis*, and Table 6 is showing *trans*-antisense gene-lncRNA pairs, and Table 7 is showing gene-lincRNAs pairs identified for *A. duranensis*. Similarly, Table 8 is showing significantly correlated *cis*-antisense gene-lncRNA pairs, Table 9 is showing *trans* antisense gene-lncRNA pairs, and Table 10 is showing gene-lincRNAs pairs identified for *A. ipaensis*. Classifications of GO terms for the highly correlated (|*CC*| ≥ 0.85) and significant (P-value < 0.05) DE gene-lncRNA pairs for both genomes are given in Table 2.

**TABLE 5.**
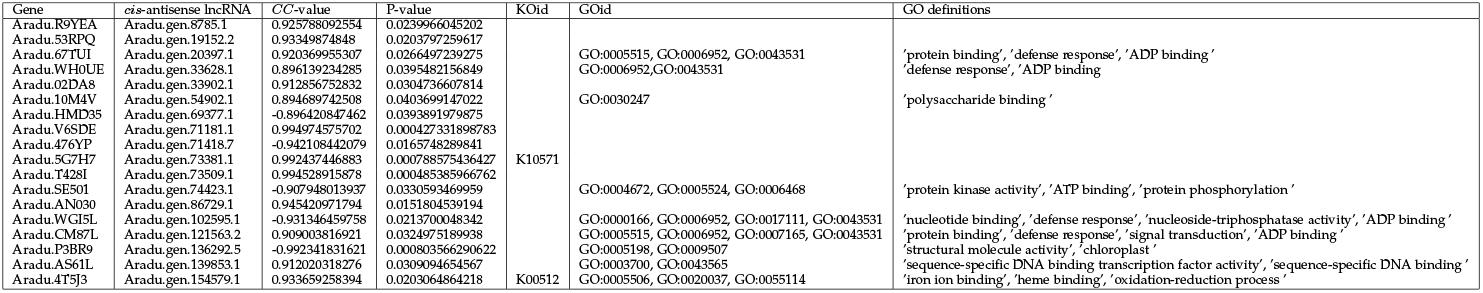
Significantly correlated DE *cis*-antisense gene-lncRNA pairs for *A. duranensis*.

**TABLE 6.**
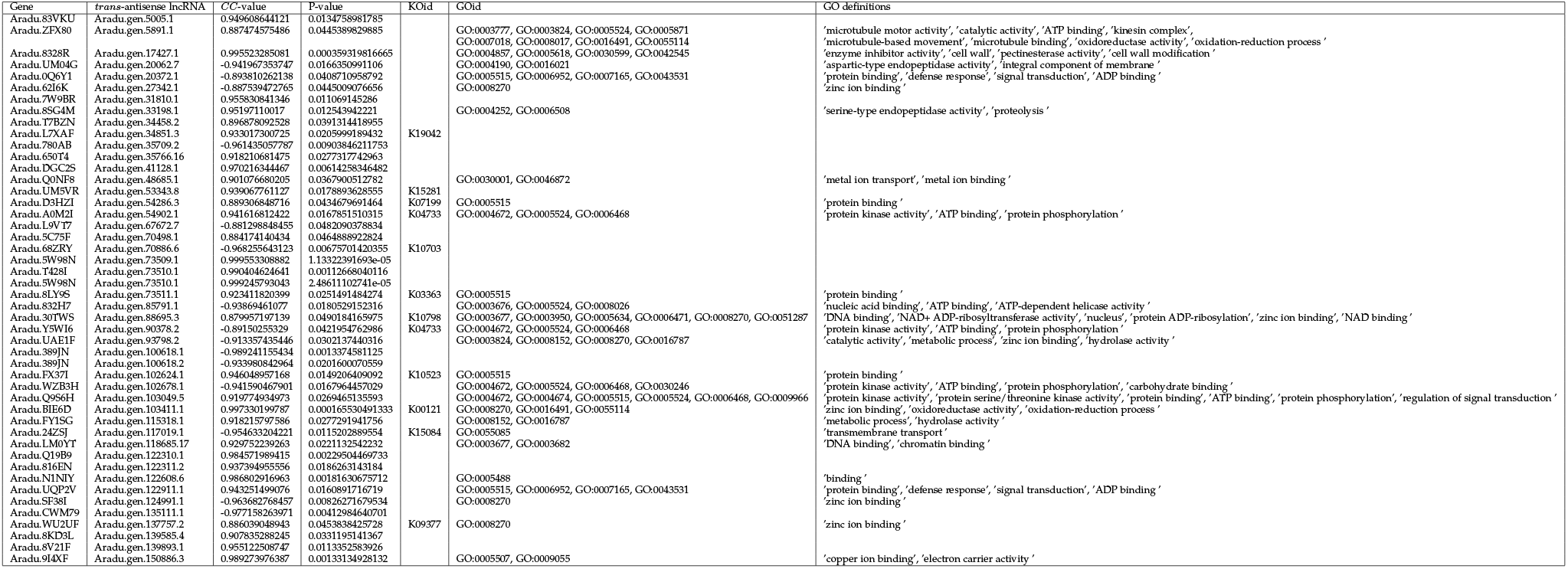
Significantly correlated DE *trans*-antisense gene-lncRNA pairs for *A. duranensis*.

**TABLE 7.**
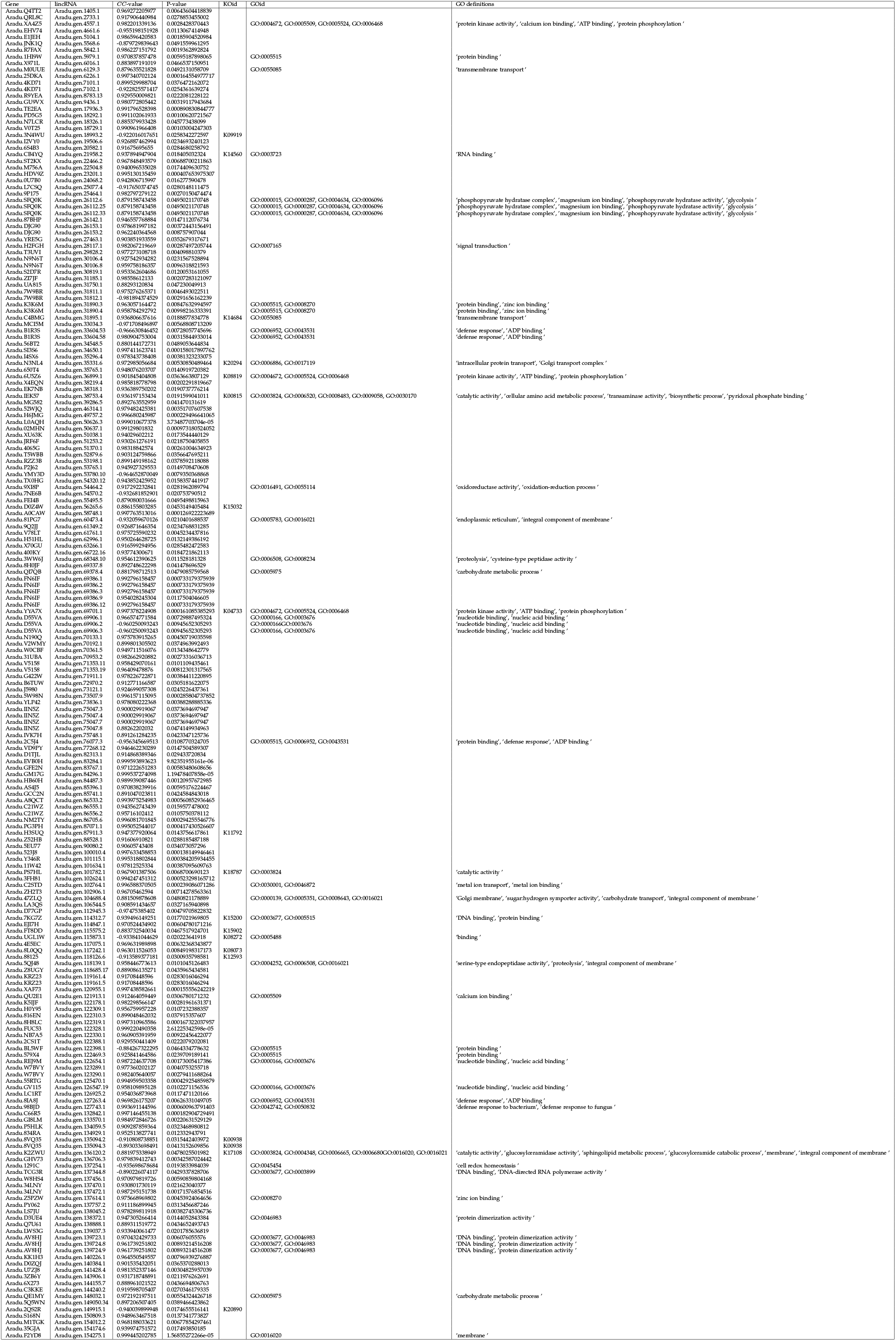
Significantly correlated DE gene-lincRNA pairs for *A. duranensis*.

**TABLE 8.**
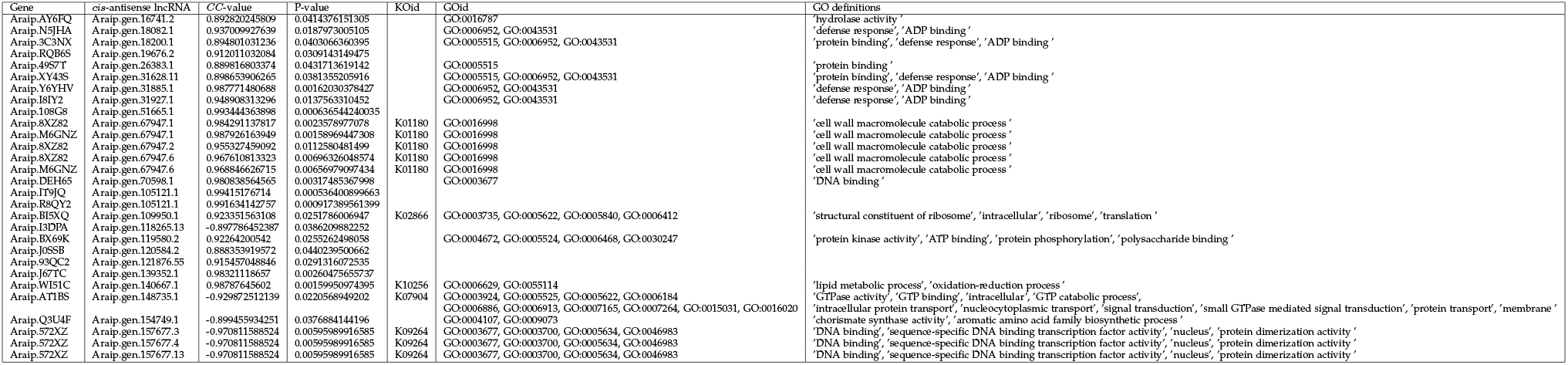
Significantly correlated DE *cis*-antisense gene-lncRNA pairs for *A. ipaensis*.

**TABLE 9.**
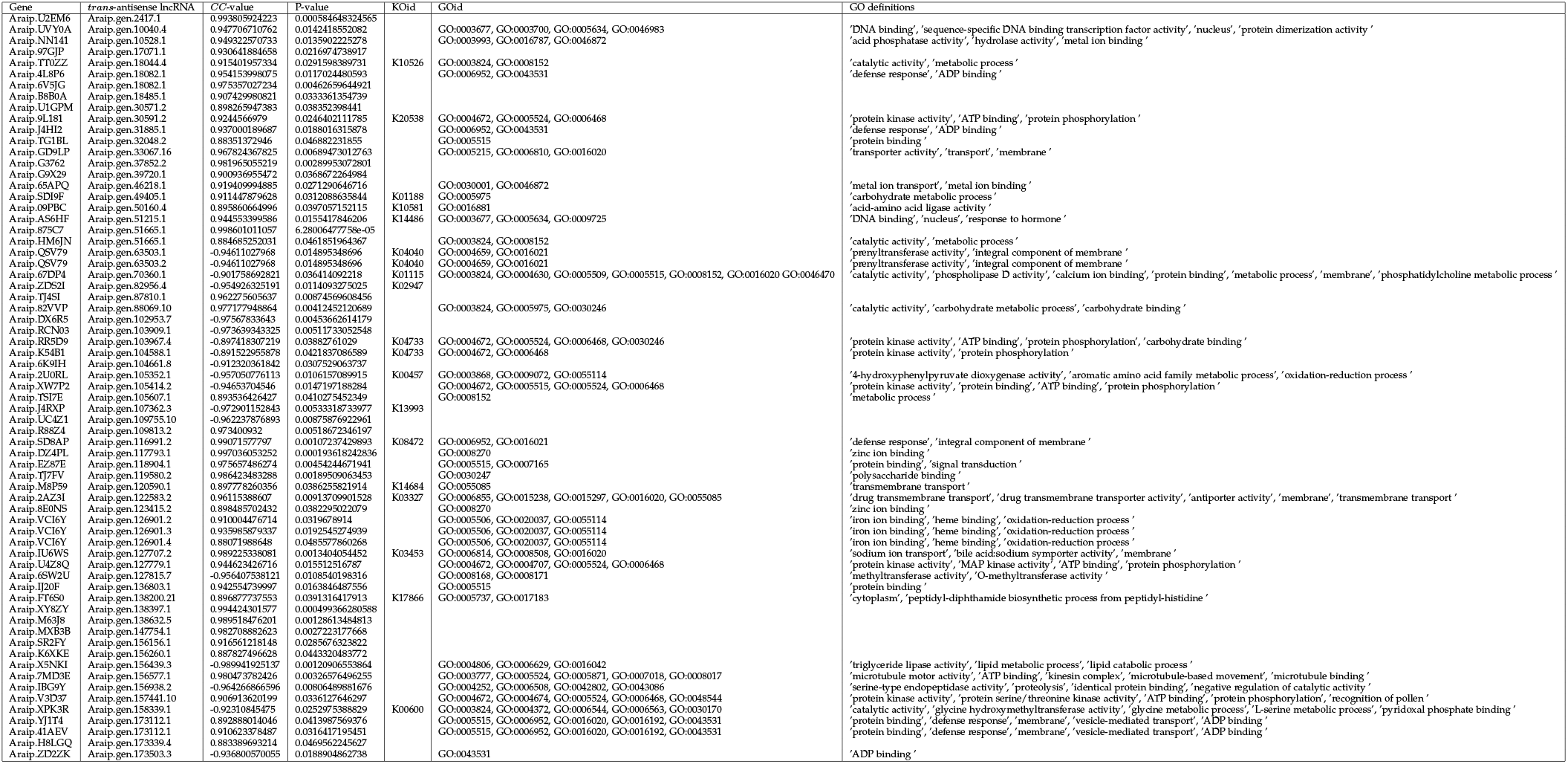
Significantly correlated DE *trans*-antisense gene-lncRNA pairs for *A. ipaensis*.

**TABLE 10.**
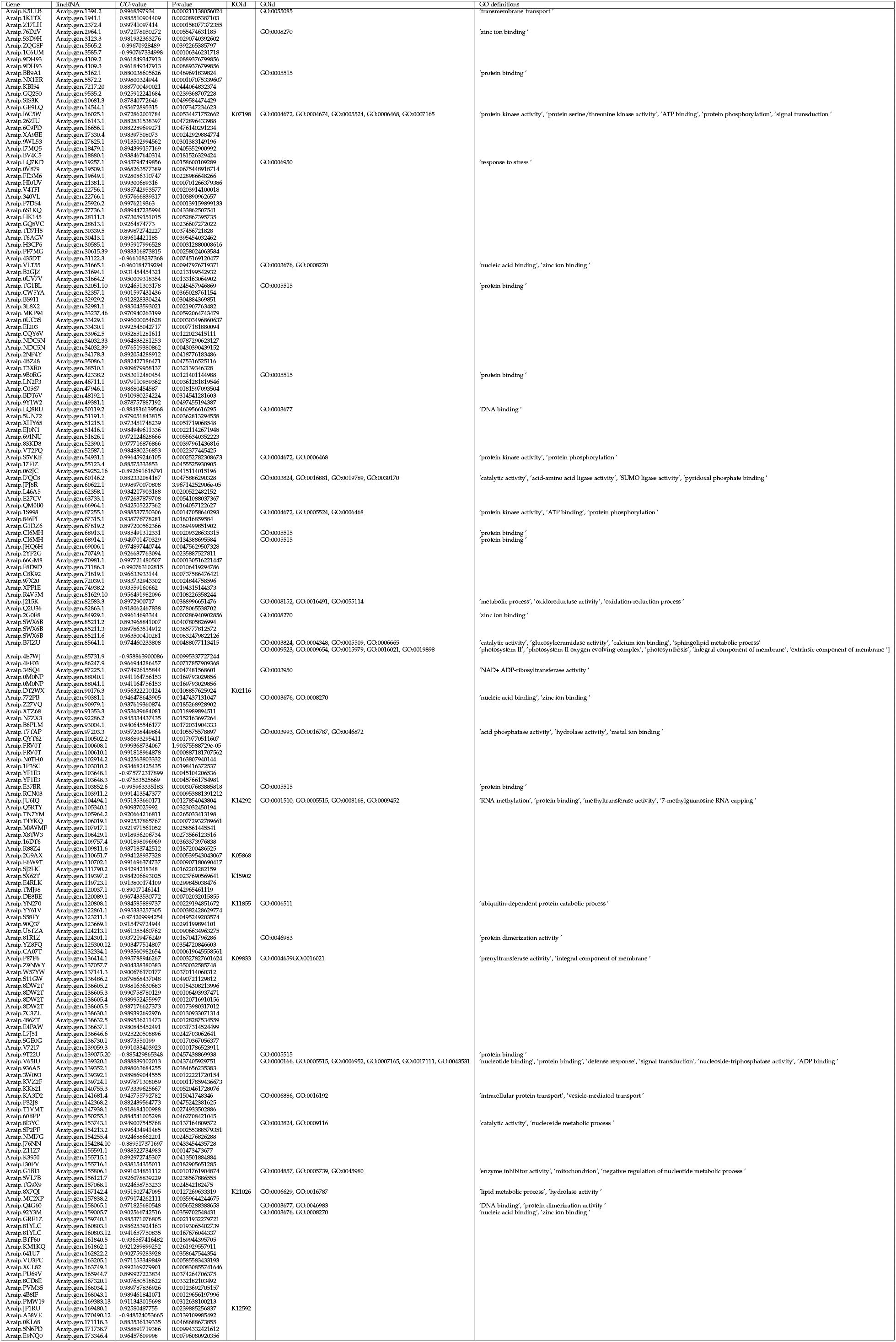
Significantly correlated DE gene-lincRNA pairs for *A. ipaensis*.

Biological pathways of the highly correlated and significant DE gene-lncRNA pairs are analyzed for the Table 5 to 10. Gene names and their respective KOids are used for the reconstruction pathway analysis (https://www.genome.jp/kegg/tool/map_pathway.html). This analysis helps to understand the implication of the major targets for lncRNAs. Biological pathways classification of the highly correlated and significant DE gene-lncRNA pairs are given in Fig. 7. About 46 metabolic networks are observed for metabolism. Where, 8 independent pathways seem to be relevant in genetic information processing and 7 signal transduction pathways. Five different cellular processes like apoptosis, cellular senescence, cell cycle are also observed. Metabolic pathways are observed for the large biological networks like Carbon, Nitrogen regulation, membrane lipid biosynthesis, and secondary metabolite biosynthesis. These are relevant with the nature of the symbiosis that involves (*i*) *C-N* exchange (*ii*) massive change in membranes for creating host-symbiont interface (*iii*) defence alert for preventing any secondary infection. The symbiont acquires photosynthetically fixed carbon in the form of dicarboxylic acids such as succinate and malate for *C-N* exchange. In turn, bacteria provide asparagine and glutamine as a nitrogen constituent to the root of *Arachis hypogaea*.

**Fig. 7.**
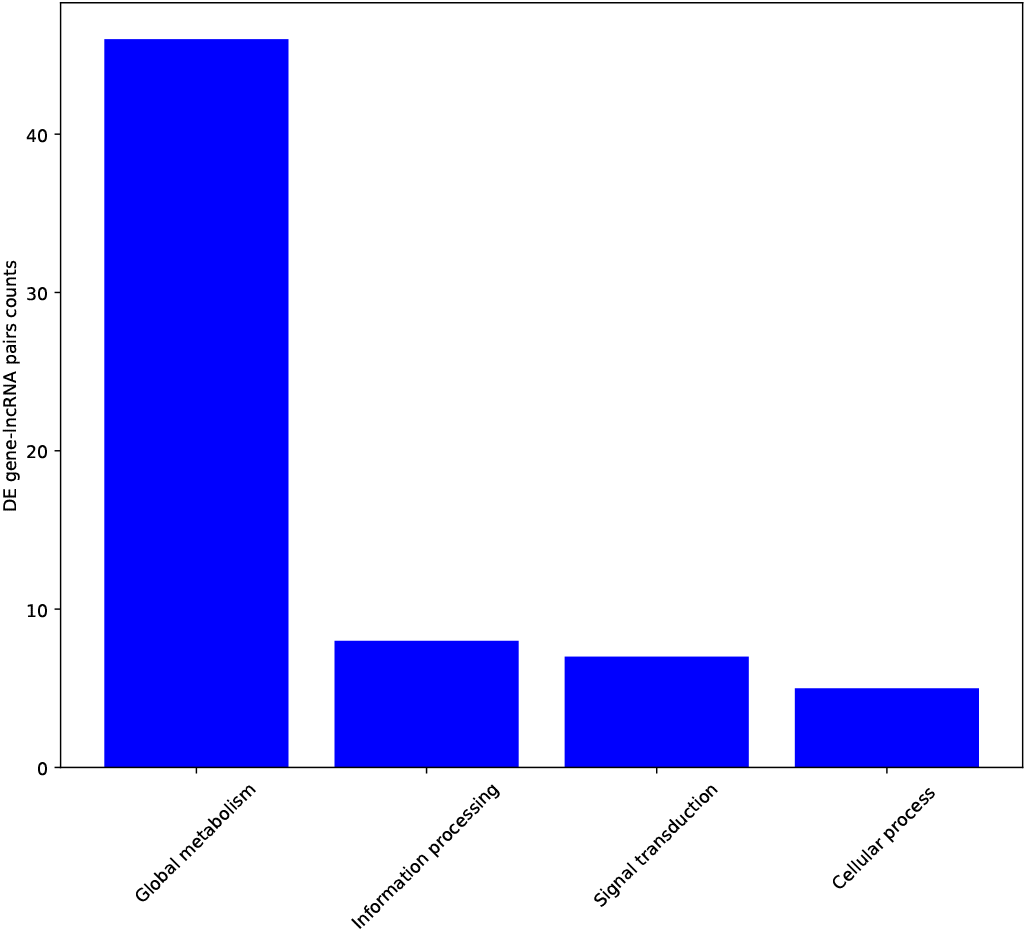
Biological classifications of pathways related to DE gene-lncRNA pairs present in Table 5 to 10.

## 4 Discussion

The relevant pathways are observed for the aspartate aminotransferase, tyrosine aminotransferase, alcohol dehydrogenase, aldehyde dehydrogenase. Aspartate aminotransferase provides aspartate by taking oxaloacetate as an amino group acceptor or glutamate by taking α-ketoglutarate, and net amination form asparagine and glutamine. Likewise, tyrosine aminotransferase provides glutamate by taking lysine as an amino group donor. These are important for the nitrogen assimilation. Other enzymes like alcohol dehydrogenase and aldehyde dehydrogenase are relevant in carbon constituent metabolism. Both of these enzymes have the significant role in glycolysis or gluconeogenesis, fatty acid degradation, and amino acid metabolisms.

In the category of membrane lipid biosynthesis enzymes like ceramide glucosyltransferase, sphingomyelin synthase, and phosphomevalonate kinase appeared to be primary targets of lncRNAs. Each of these enzymes has the significant role in membrane microdomain formation which is a prime requisite for assembling the signalosome to initiate the symbiosis in the epidermal layers of the root. Implementation of the symbiosis is established by the nutrient exchange in the developed root nodule. In the category of secondary metabolites, an important target is geranyl-geranyl diphosphate transferase which is an evolutionarily conserved class of enzymes in Archaea, Bacteria, and Eukarya. This participates in a broad range of biosynthetic pathways including cholesterol, porphyrin, carotenoids, ubiquinone, and iso-prenoids. Accordingly, Homogentisate phytyltranseferase which has a role in tocopherol biosynthesis, ubiquinone, and terpenoid-quinone biosynthesis pathway is also regulated by lncRNAs during symbiosis. Another identified target of lncRNAs during symbiosis is acetylajmaline esterase of Indole alkaloid biosynthesis pathway. This suggests a tight regulation of alkaloid synthesis during beneficial plant-microbe interactions. Together, strict regulation and generation small molecule barcode seems to be a guiding principle for a beneficial interaction like root nodule symbiosis.

## 5 Conclusion

Peanut *A. hypogaea* genome has several lncRNAs which have epigenetic controls during root nodulation process. Expression patterns of lncRNA and genes are correlated to check these epigenetic controls. Highly correlated and significant DE gene-lncRNA pairs are listed, and their biological significance are explained here. This study is also the first explanation of gene-lncRNAs pairs of peanut genome which creates a roadmap for the further epigenetic research.

